# A hymenopteran odorant alerts flies to bury eggs

**DOI:** 10.1101/2021.09.30.462443

**Authors:** Shaun M. Davis, Gregory T. Chism, Megan M. Maurer, Julio E. Trejo, Ricardo J. Garcia, Todd A. Schlenke

## Abstract

Ants are ubiquitous and consume insects at all life stages, presumably creating a strong selective pressure for ant avoidance behaviors across insects. The insect egg stage can be especially defenseless against predation given that eggs are usually immobile and unguarded, suggesting insect mothers may have evolved oviposition strategies to minimize the ant predation risk to their offspring. Given the lack of parental care in most insects, these oviposition strategies would likely be innate rather than learned, since insect mothers are not usually present to assess predation of their eggs. Here, we use the vinegar fly *Drosophila melanogaster* as a model system for examining parental defensive responses to ant presence. Flies usually lay eggs partially inserted into the food substrate, although some are laid on top of the food and a few are inserted deeply into the food. We found that exposure to ants significantly alters fly oviposition depth: the proportion of eggs on the food surface decreased while the proportion of buried eggs increased. Buried eggs survive ant foraging bouts better than surface eggs, showing that this oviposition depth behavior is adaptive. This induced behavior is conserved across the genus Drosophila and is dependent on the fly olfactory system: anosmic mutant flies fail to bury their eggs in the presence of ants, and ant odor extracts are sufficient to induce egg burying. To further delineate the ant lineages to which flies respond, we exposed flies to the odors from numerous species of ants and other insects. Surprisingly, flies buried their eggs in response to the odors of nearly all hymenopterans tested, including hymenopteran groups that flies rarely interact with in nature like bees and paper wasps. Our data suggest that hymenopterans possess a conserved and ancient odorant, and that drosophilids evolved a mechanism for sensing this odorant early in their evolution as a means of protecting their offspring from ant predation. This study sheds light on the ecology and mechanisms underlying a common biotic interaction in nature, that between insect parents ands the ants that would consume their offspring.

## Introduction

One of the greatest threats that animals face in nature is predation – being killed and eaten by another organism (Bowman & Hacker, 2020). A common type of defense against predation are defensive behaviors, e.g. where animals hide or escape from nearby predators. Organisms have also evolved to assess and behaviorally respond to the mere risk of predation (Hermann & Landis, 2017), and will even engage in behaviors that protect their close relatives (e.g. offspring) from predation. Because predation results in death, antipredation behaviors, especially in non-social organisms, are usually innate rather than learned, encoded in an organism’s germ line and brain (Baker et al., 2001; Ren & Tao, 2020). The neurogenetic basis of how naïve prey organisms perceive and respond to predators to which they have never been exposed remains poorly understood. Therefore, it is useful to develop a model system for studying the mechanistic basis of innate antipredation behaviors.

The vinegar fly *Drosophila melanogaster* has been a genetic ‘model organism’ for more than a century, and has already proven useful for understanding interactions with a different kind of biotic threat: parasitism (Lemaitre & Hoffmann, 2007). Furthermore, recent technological innovations have made it possible to genetically manipulate the activity of relatively small groups of neurons in the fly nervous system, and thus define the neurological circuitry underlying defensive behaviors (Venken et al., 2011). These tools have been used to uncover the neurological basis of fly responses to simulated predation scenarios, such as a visual ‘looming’ stimulus (Ache et al., 2019; Morimoto et al., 2020; Muijres et al., 2014). Nevertheless, experiments with natural predators will yield a more nuanced and complete picture of fly defense behaviors. Because flies and their potential predators are relatively small, these natural predator-prey interactions can be studied in controlled lab settings. However, one drawback of developing flies as a model for anti-predation behaviors is that natural history data describing the relative importance of different predator types is limited for natural populations of *Drosophila* (Markow, 2015; Reaume & Sokolowski, 2006; Soto-Yéber et al., 2018).

A large diversity of generalist predators likely consume *D. melanogaster* in nature. We have observed adults being caught in the air or picked off surfaces by predatory flies, dragonflies, spiders, lizards, and hummingbirds. Flies have evolved constitutively erratic flight patterns, as well as induced behaviors like evasive flight maneuvers, avoidance, freezing, jumping, and posturing to escape capture in these contexts (Combes et al., 2012; de la Flor et al., 2017; Muijres et al., 2014; Parigi et al., 2019). Less is known about predation of *D. melanogaster* eggs, larvae, and pupae in nature, although we have observed rove beetles, predatory beetle larvae, predatory fly larvae, and ants consuming these juvenile stages. Given that fly eggs and pupae are immobile and that larvae are relatively slow crawlers, options for behavioral defenses against predators in these life stages appear limited. However, some constitutive behaviors have presumably evolved to limit predation of juvenile flies, such as adult females preferring to oviposit in food crevices and chemically masking their eggs with pheromones, larvae preferring darker (more hidden) parts of the food and often burrowing as they are eating, and pupating larvae dispersing away from the food and gluing themselves to a substrate (Borne et al., 2021; Narasimha et al., 2019; Rockwell & Grossfield, 1978; Sawin-McCormack et al., 1995; Soto-Yéber et al., 2018; Vijendravarma et al., 2013)

Female oviposition choices are particularly interesting because they can represent trans generational anti-predation behaviors (a type of parental care) (Refsnider & Janzen, 2010). The decision about where, when, and how to lay an egg is complicated and relies on multiple kinds of sensory information (Cury et al., 2019; Richmond & Gerking, 1979; Rockwell & Grossfield, 1978; Wang et al., 2020; Zhang et al., 2020). This information includes the mating and reproductive status of the female, aspects of the abiotic environment like the weather, the time of day and season, aspects of the food source like water content, stiffness, color, odor, and taste, and aspects of the biotic environment like presence of conspecifics and potentially the presence of biotic threats. Might flies alter their oviposition choices (i.e. trigger an innate induced behavior) in the presence of predatory threats to their offspring?

Ants are ubiquitous and occupy diverse niches in nearly all natural ecosystems (Hölldobler & Wilson, 1990). Many are facultative or obligate predators of other organisms, including insects of all life stages (Fernandes et al., 2012), which provide necessary proteins and fats for colony growth (Hölldobler & Wilson, 1990). A number of studies have shown that presence of ants, and specifically visual or olfactory ant signals, cause diverse insect species to avoid oviposition in the ant-infested area (Freitas & Oliveira, 1996; Sendoya et al., 2009; Taylor et al., 1998; Van Mele et al., 2009). Although published data on interactions between *D. melanogaster* and ants are scarce (Soto-Yéber et al., 2018), ants have been shown to be important predators of the eggs and larvae of other *Drosophila* species (Escalante & Benado, 1990; Lewis & Worthen, 1992; Worthen et al., 1993). Anecdotally, we have noticed that rotting fruit traps meant to attract *D. melanogaster* attract far fewer flies when ants are present, and we have seen ants carrying off what appear to be *D. melanogaster* eggs and larvae.

Here we tested whether naïve flies alter their oviposition behavior in the presence of predatory ants. We discovered that flies push their eggs deeply into the food substrate when exposed to ants, which protects eggs from ant predation. This innate, induced oviposition depth behavior is a conserved trait across the genus Drosophila. Unlike previous work with other fly predators (de la Flor et al., 2017), olfaction, but not vision, is required for the oviposition depth behavior. Furthermore, we show that flies deploy this behavior in response to diverse ant species, as well as other hymenopterans. This system can serve as a model for how innate threat recognition and downstream defensive behaviors are encoded in the germline and hardwired into the brain.

## Materials and Methods

### Species used

#### Flies

All flies were raised on cornmeal molasses food (10L water, 75g agar, 275g yeast, 520g cornmeal, 110g sugar, 1046g molasses, 45mL propionic acid, and 100mL 20% (w/v) Tegosept) and maintained at 24°C in ∼60% humidity with a 16:8 light:dark cycle. We used a *D. melanogaster* Oregon R (OreR) strain as our wild-type strain. The following *D. melanogaster* mutants were obtained from the Bloomington Drosophila Stock Center (with stock number given): vision mutants GMR-hid (5771) and ninaB^1^ (24776), olfaction mutant orco^2^ (23130), the driver strain Or49a-GAL4 (9985), and the responder strains UAS-Kir2.1 (6595) and UAS-hid (65403). The mutant chromosomes from these strains were crossed into the OreR background to reduce genetic variability. White-eyed w^1118^ flies were provided by Daniela Zarnescu and used as hosts for growing parasitoid wasps. Of the other *Drosophila* species used, wild *D. simulans* were caught in Tucson, AZ in 2018 and maintained in the lab as a single strain. *D. yakuba* and *D. virilis* were acquired from the Drosophila Species Stock Center (stock numbers 14021-0261.01 and 15010-1051.87, respectively).

#### Ants

A laboratory culture of *Pheidole hyatti* was originally started in June 2018 from a single queen collected post-nuptial flight and kept in a small 5ml cotton-stopped tube. Once the queen’s first brood emerged the housing tube was placed into 17.5cm x 12.5cm x 6cm plastic container with inside walls coated in ‘insect-a-slip’ (BioQuip product #2871A) to prevent escape. The colony was given cotton ball stopped, water-filled 5 ml plastic tubes, and were fed ad libitum weekly with both a 2 ml microcentrifuge tubes of honey water (1/4 teaspoon per 50 mL water), and 1/8 of a fresh-frozen cockroach (approximately 0.075g) (*Nauphoeta cinerea*). During acclimation, the colony was kept in a laboratory at 20°C with a 12:12 h light cycle and 20-25% relative humidity. Once the colony outgrew the initial nest chamber, larger 3.5 cm diameter water-filled glass tubes, stopped with cotton, and a housing container measuring 31.5cm x 23.5cm x 10.5cm was provided. Other species of wild ants were trapped around Tucson, AZ using small amounts of protein (tuna) and sugar (honey) arranged on pieces of cardboard placed near active ant trails. After roughly 1 hour, ant-covered cardboards were transferred to gallon-sized zip lock bags and brought back to the lab, where most were frozen at -20°C for a later experiment (Fig. 5). Because the laboratory *P. hyatti* colony collapsed during the course of these experiments, we turned to readily-available *Forelius mccooki* wild ants for many live exposure experiments. Live *F. mccooki* workers were collected and placed in a 31.5cm x 23.5cm x 10.5cm bin (walls coated with insect-a-slip) and provided with 2 cm and 3.5 cm diameter cotton-stopped glass tubes of water. Live ants were never reused across experiments.

Ant species were identified morphologically to genus using a dichotomous key (Fisher & Cover, 2007), and identified to species by regional species level keys (see Supplementary Table 1 for all keys and sources). No species used in this study are protected or endangered. To confirm morphological identifications, DNA was extracted from the ant samples for Sanger sequencing. First, 10-50 ants were homogenized in 20-30 μL extraction buffer (0.1M Tris-HCL, pH 9.0, 0.1M EDTA, 1% SDS) with a Kontes pellet pestle in a 1.5 mL Eppendorf tube. Additional extraction buffer was added to bring the volume up to 500 μL, then the samples were incubated for 30 min in a 70°C water bath. Cellular material was precipitated and pelleted by adding 70 μL of 8M potassium acetate to each sample, incubating on ice for 30 min, and centrifuging at 13000 rpm for 15 min at 4°C. The supernatant was transferred to a fresh Eppendorf tube, then the DNA was precipitated and pelleted by adding 300 μL isopropanol and centrifuging at 13000 rpm for 5 min at room temperature. After a second clean-up spin, the resulting DNA pellet was washed with 1 mL 70% ethanol and centrifuged at 13000 rpm for 5 min at room temperature to re-pellet. The pellet was allowed to air dry for 1-2 minutes, then resuspended in 75 μL nuclease-free water and stored at -20°C. The mitochondrial *cytochrome oxidase I* (*COI*) gene was sequenced to identify the different ant species. Following the GoTaq Green Master Mix (Promega, WI, USA) protocol, we generated 25μL reactions containing 12.5 μL GoTaq, 7.5 μL nuclease-free water, 2 μL of both LCO1490 and HC02198 primers (Folmer et al., 1994), and 1 μL extracted DNA. Polymerase chain reaction (PCR) conditions were as follows: 94°C for 5 min, 37 cycles of 94°C for 1 min, 51°C for 1 min, 72°C for 1 min, followed by 72°C for 2 min and a hold at 4°C. The PCR products were visualized on a 1% agarose gel with SYBR safe DNA gel stain (Thermo Fisher Scientific) to confirm single DNA bands were present. PCR products were purified using the QIAquick PCR Purification Kit (Qiagen) and then sequenced at the University of Arizona Genetics Core using the PCR primers. All sequences can be found in Supplementary File 1.

#### Wasps

The *Drosophila* parasitoid wasp *Leptopilina heterotoma* (strain Lh14) and *L. boulardi* (strain Lb17) (Schlenke et al., 2007) were used in various wasp exposure assays, and a number of other live parasitoid species maintained in the Schlenke lab were used in a later experiment (Fig. 5). To culture most of these wasps, adult *D. melanogaster w*^*1118*^ flies were allowed to lay eggs in vials containing *Drosophila* media for 3-4 days, after which the flies were replaced with 5-10 female and 2-3 male wasps. The wasps *L. clavipes, Ganaspis brasiliensis*, and *Asobara tabida* were reared the same way, except that they were grown on the host species *D. virilis* and in bottles rather than vials. The adult wasps were aged for at least 2 days before being used in exposure experiments. Live wasps were never reused across experiments.

#### Other species

A variety of other arthropod species were used in an experiment to determine the range of organisms to which *D. melanogaster* responds (Fig. 5). Many of these species were collected around Tucson, AZ, and identified by morphology to the most specific taxonomic group possible. Several other species were provided by research labs at the University of Arizona. See Supplementary Table 2 for more information about all the species used. The NCBI taxonomy ID was used to create a phylogeny of the arthropods used using phyloT and iTOL software (Letunic & Bork, 2021).

### Exposure experiments

Flies were grown in 6oz square bottom culture bottles (Genessee Scientific). On day 0, all adults were cleared from the bottles, the flies were allowed to eclose for three days, then collected and kept in standard molasses food vials for roughly 24 hours until day 4. New vials were prepared with 0.5g ± 1mg Instant Drosophila Medium (Carolina Biological Supply Company) and hydrated with 1.6 mL water with 1% red food coloring (McCormick) (to enhance contrast between the food and eggs). The vials were supplemented with 3-4 drops of hydrated yeast to promote egg development in female flies. The flies were sorted into groups of ten females (of the appropriate genotype) plus two OreR males per vial and kept overnight on this food until day 5 to allow them to recover from CO_2_ exposure before the experiment. The experimental vials were also prepared using the same red instant food but without the added yeast. For live insect exposure trials, on the day prior to the start of the experiment eight live insects, either *F. mccooki* or *P. hyatti* ants, or Lh14 or Lb17 wasps, were added to the experimental vials to allow the odorous compounds to accumulate (day 4). Control, unexposed vials contained no insects. The following day (day 5), the 2-5 day old flies were flipped to the experimental vials, with or without live insects or insect odors. For the whole-body wash odor exposure trials (see below), 50 μL of the solvent (control) or odor extract was added to each experimental vial. The vials were kept in a fume hood to allow the solvent to evaporate for fifteen minutes before the flies were added to start the experiment.

For all exposure experiments, we ran five replicates per treatment. The flies were kept in the experimental vials for 24 hours at 24°C in ∼60% humidity with a 16:8 light:dark cycle, then all insects were removed and the vials were moved to -20°C to stop egg development. The following parameters were used to categorize each egg position: ‘surface’ eggs were those for which their entire circumference, from anterior to posterior ends, was entirely visible; ‘buried’ eggs were those where the position at which the dorsal appendages emerges from the eggshell was located below the surface of the substrate; ‘partial’ eggs were those that were positioned in the food substrate, but not to the extent of the buried eggs. Unlike *D. melanogaster* eggs, *D. virilis* eggs have two pairs of dorsal appendages, so the more posterior pair was used to determine the ‘buried’ position. To calculate the egg position index (EPI), we used the following equation: EPI = (S – B)/T where S is the number of eggs on the surface, B is the number of eggs that are buried, and T is the total number of eggs across all three categories.

### Odor extracts

All insects were frozen at -20°C for at least 24 hours prior to odor extraction. A freeze-thaw cycle was shown to improve detection of ant volatile compounds (Chen, 2017). We used 1.5 mL Eppendorf tubes, washed two times with 500 μL solvent, for the odor extraction. Initial experiments with ants or parasitoid wasps used 70 insects washed with 350 μL solvent. Tubes were vortexed for 1-2 minutes, centrifuged briefly, then 50 μL aliquots (representing the odor equivalent of ten insects) were pipetted out of the tube and added to each of the five replicate experimental vials. For control vials, the pure solvent underwent the same procedure but without insects. For odor extractions from especially small or large arthropod species, we used body mass instead of number of individuals to determine the solvent-to-insect ratio. A minimum of 30 mg was washed with 350 μL solvent. See Supplementary Table 1 for more information about how body washes were generated.

### Ant predation experiment

OreR flies laid eggs on standard molasses food in 5.5cm diameter petri dishes for up to two hours. Eggs were gently removed and manually positioned on 5.5cm diameter petri dish filled with 0.5% agarose gel with red food coloring. Surface eggs were set on top of the agarose gel, partially buried eggs were inserted lengthwise roughly 50% into the gel, and the buried eggs were inserted until the gel covered the point at which the dorsal appendages connect to the eggshell (twenty eggs per category for a total of sixty eggs). The *P. hyatti* laboratory colony was used under the assumption that it displays normal foraging activity. A secondary 17.5cm x 12.5cm x 6cm plastic container, with walls coated in ‘insect-a-slip,’ was connected to the main colony chamber by a 2cm (length) x 1.5cm (diameter) plastic tube plugged with cotton. The cotton separator was then removed, and ants were allowed to freely explore the smaller chamber for thirty minutes before the dish with fly eggs was added. The ants were allowed to forage for thirty minutes before removal of the egg dish, at which point the remaining eggs were counted. The connection between the colony chamber and the foraging chamber was then blocked, the remaining ants were transferred back to the colony chamber, and the bottom of the foraging chamber was washed with ethanol to remove any ant pheromone trails. After one hour, the cotton separator was removed and the next replicate trial began. Two replicates were run per day over the course of ten days, for a total of twenty replicates. After counting the remaining fly eggs, we scored egg hatching rate 48 hours after the ant exposure to confirm that the remaining eggs were viable.

### Testing tradeoffs to deep oviposition

To test whether buried fly eggs show lower survival or slower development to the adult stage, we had OreR flies lay eggs on standard molasses food in 5.5cm diameter petri dishes for two hours. Groups of thirty eggs were each transferred to a vial with molasses food and manually positioned such that all eggs were either on the surface or buried deeply in the food. Starting eleven days post egg lay, the number of flies that eclosed per vial was counted daily. To test whether buried fly eggs are more susceptible to a biotic risk (being further buried by foraging fly larvae), twenty white-eyed (w^1118^) third instar larvae were added to the vials of fly eggs, and eclosion was assayed. To test for cold temperature risk, vials with fly eggs were placed overnight (∼15 hours) at 4°C, then transferred to 24°C to allow the flies to continue development, before assaying eclosion. To test for risk of drowning, vials with fly eggs were sprayed with water using a 4oz fine mist sprayer bottle such that the bottom of the vial had a thin layer of water, before assaying eclosion.

To test whether there is a tradeoff to adult female flies when ovipositing deeply into the food, we measured the duration of the oviposition phase and the clean-and-rest phase post-oviposition by taking videos of groups of ovipositing flies. Batches of five OreR females plus one male were collected from bottles three days after being cleared and kept on molasses food overnight in rectangular 3 cm (length) x 3 cm (width) x 6.3 cm (height) chambers. The following day, five female wasps (strain Lh14) were added and kept co-housed with the flies overnight. The next day, all insects were transferred to a new rectangular chamber with molasses food. Two strips of parafilm were placed in parallel on top of the food, leaving a 2-3 mm wide area available for oviposition at a depth of field amenable to video capture. Multiple 1-hour videos were recorded per day using a Basler 2.3-megapixel acA1920-155μm camera (Graftek Imaging). Frames were captured using Pylon Viewer (version 6.2.4.9387 64-bit) at 15 Hz, and converted to .avi video format using MATLAB code (Chowdhury et al., 2021). Egg position was scored, then the oviposition and clean-and-rest phase durations were determined by examination of the video files.

### Statistical analysis

All graphs and statistical tests were performed using GraphPad Prism 9 (v 9.2.0) software. All data were tested for normality using the Shapiro-Wilk test. Oviposition rates, egg position index, and eclosion rates were tested using the unpaired, two-tailed t-test with Welch’s correction or two-way ANOVA followed by Dunnett’s multiple comparison test against the control condition. The Brown-Forsythe and Welch’s ANOVA with Dunnett’s multiple comparison test was used on egg position data with multiple, independent samples. Fly egg survival after ant predation, oviposition timing, and clean-and-rest phase timing were compared using the nonparametric Kruskal-Wallis test with Dunn’s multiple comparison test. Throughout the paper, statistical notation are as follows: * p<0.05, **p<0.01, ***p<0.001, ****p<0.0001, ns: not significant.

## Results

### Flies bury their eggs when exposed to ants

To test whether adult female flies change their oviposition behavior in the presence of predatory ants, we co-housed flies with either wild-caught *Forelius mccooki* or laboratory raised *Pheidole hyatti* ants for 24 hours. Unlike with exposure to parasitoid wasps (Supp. Fig. 1A and C) (Kacsoh et al., 2015; Lefèvre et al., 2012; Lynch et al., 2016), female flies did not reduce oviposition in the presence of these ants (Fig. 1A and B). The non-significant trend towards reduced egg numbers from the *P. hyatti* exposure is likely due to the ants actively capturing the adult flies during the experiment, as exemplified by flies that had sections missing from their wings or legs at the end of the experimental period (Fig. 1C). Ants of the genus *Pheidole* have been described as highly aggressive in terms of prey capture (Traniello, 2010).

**Fig. 1:**
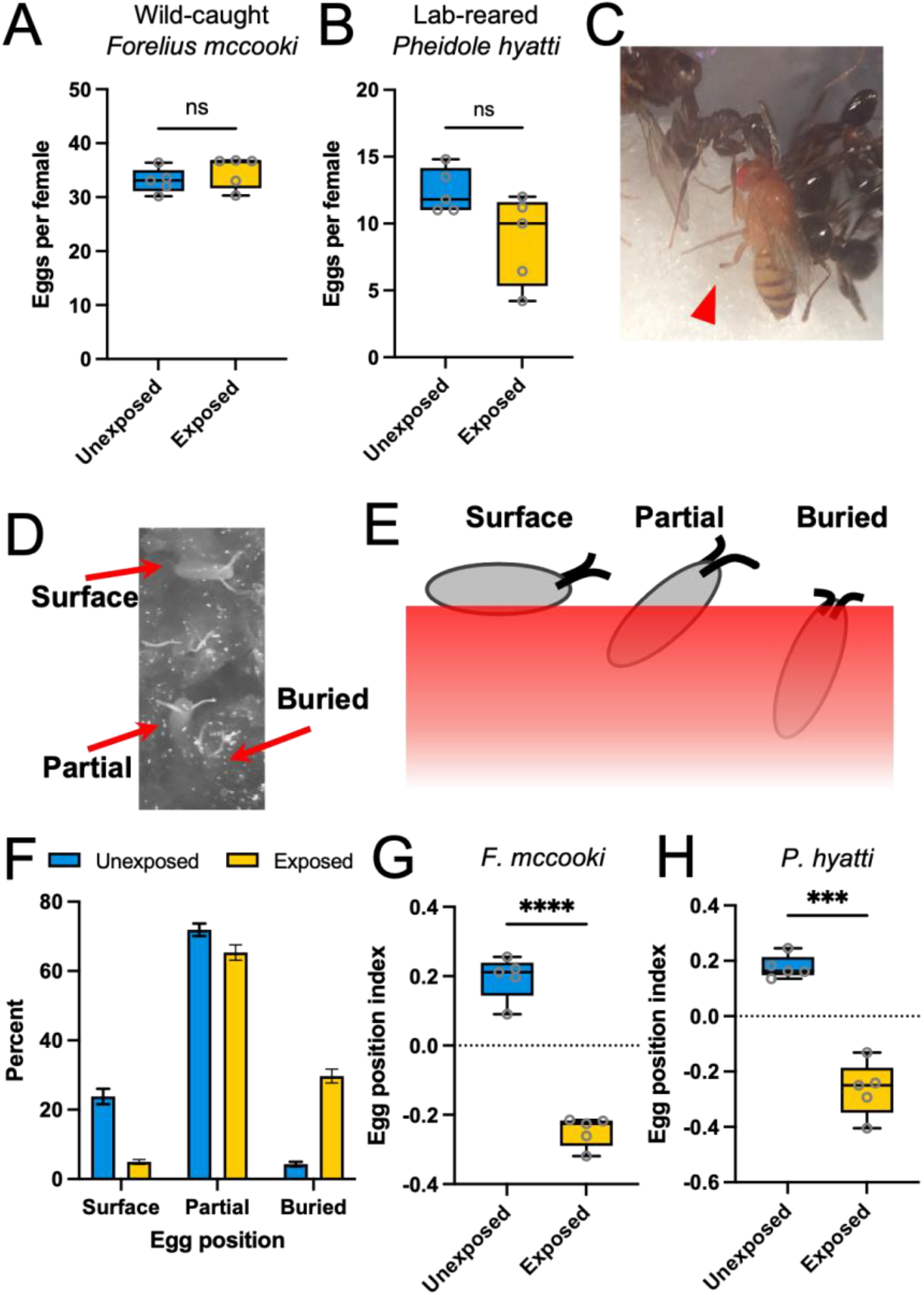
Flies alter egg depth when exposed to ants. *D. melanogaster* flies exposed to either wild-caught *Forelius mccooki* (**A**) or lab-reared *Pheidole hyatti* (**B**) ants did not reduce oviposition rates (unpaired, Welch’s t-test; *F. mccooki*: *P*=0.3551; *P. hyatti*: *P*=0.0713). The lower, but non-significant, exposed egg numbers in (B) are likely due to some flies being caught by the ants (**C**). The arrowhead indicates a leg that was severed. We did not observe any flies being caught by *F. mccooki* ants. (**D**) Photo of eggs in an exposure experiment, indicating the three different egg depth categories. (**E**) Schematic of oviposition depth categories. (**F**) Ant exposure significantly alters the distribution of eggs in each category (χ^2^=28.93, *P*<0.0001, bars represent mean ± S.E.M). (**G** and **H**) Both *F. mccooki* and *P. hyatti* ants significantly altered the egg position index towards a preference for buried eggs rather than surface eggs (unpaired, Welch’s t-test: *F. mccooki*: *P*<0.0001; *P. hyatti*: *P*=0.0002).

Although there was no reduced oviposition, we noticed while counting the fly eggs that the eggs of ant-exposed flies tended to be less visible than those of control unexposed flies. Flies oviposit their eggs either fully on the surface of the food substrate, partially inserted into the food, or completely submerged beneath the surface (Fig. 1D-E). Further examination revealed that the majority of the eggs, regardless of ant exposure condition, were laid partially submerged into the food substrate (Fig. 1F), but that the second most common category switched depending upon ant exposure. Unexposed flies laid more of their eggs on the surface while those exposed to ants pushed more of their eggs deeply into the food substrate (Fig 1F). To better describe this alteration in oviposition depth, we calculated the egg position index, which is the proportional difference between the surface vs buried eggs (see methods). A positive value indicates a stronger preference to lay eggs on the surface of the food while a negative value indicates a stronger preference for burying eggs under the food. Exposure to both *F. mccooki* and *P. hyatti* ants significantly altered the flies’ oviposition preference towards the buried egg category (Fig. 1G-H). Interestingly, this behavioral modification also occurs when flies were exposed to parasitoid wasps (Supp. Fig. 1B and D).

### Buried eggs survive ant predation

We hypothesized that eggs laid deeper into the food substrate would be protected against ant consumption. We provided the laboratory culture of *P. hyatti* ants with fly eggs manually positioned at three different depths, similar to the three natural depth categories (Fig. 1D-E, 2A). After a 30 minute ant foraging bout, the remaining fly eggs were counted. Eggs positioned on the surface of the media were the most susceptible to predation, while eggs positioned beneath the surface were strongly protected (Fig. 2B). Indeed, ants readily removed the eggs on the surface but struggled (attempted and often failed) to remove the deeply positioned eggs (Supp. videos 1 and 2). Of the eggs remaining after the ant foraging bout, hatching rates were high and similar across the depth categories (Fig. 2C), showing that the ants were not consuming or otherwise harming the buried eggs on-site. These results suggest that the fly oviposition depth behavior is an adaptation to protect fly offspring from ant predation.

**Fig. 2:**
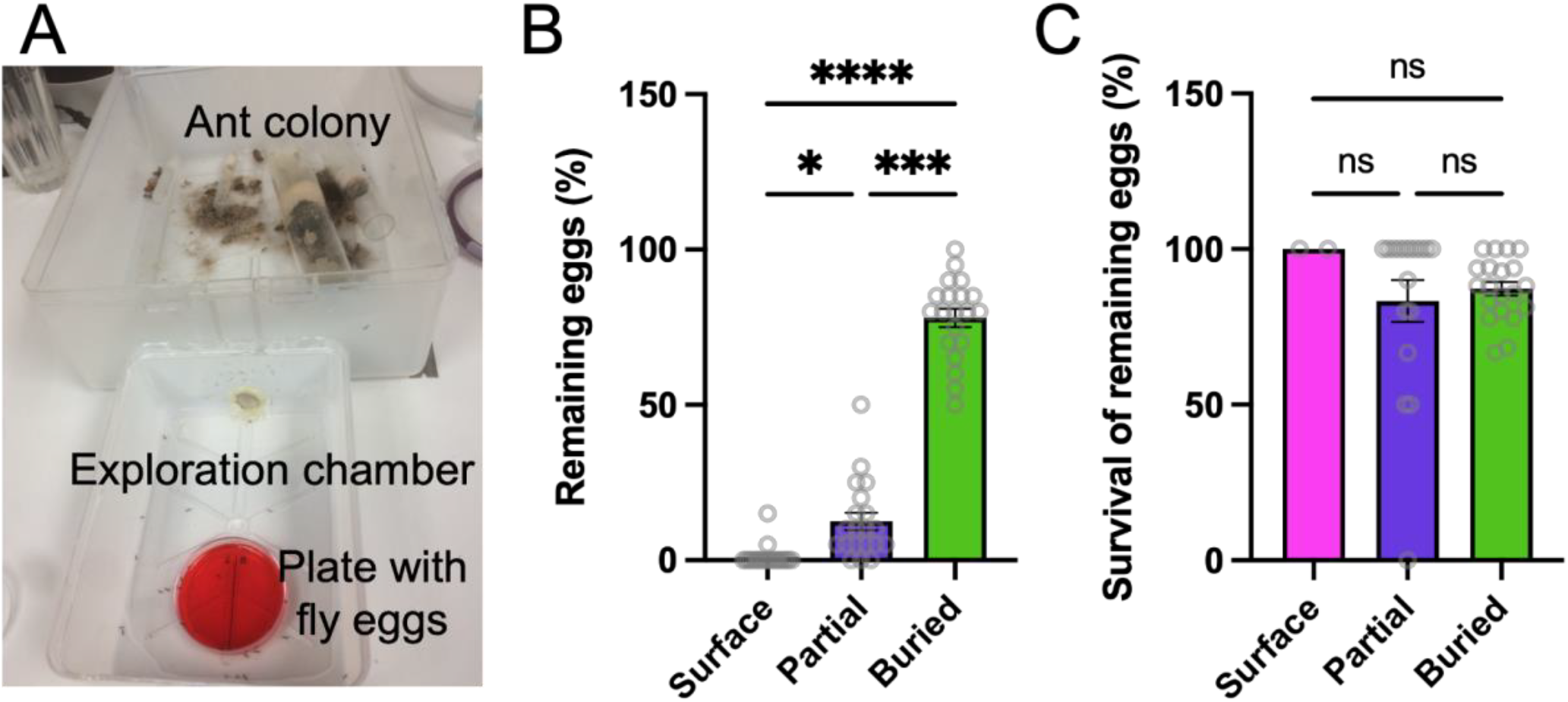
Fly eggs positioned deeply are protected from foraging ants. (**A**) Photo of ant foraging experiment with *P. hyatti* ants. (**B**) Buried fly eggs were significantly more likely to remain after ant foraging, while surface and partially buried eggs were removed by the ants (Surface vs. Partial: *P*=0.0135; Surface vs Buried: *P*<0.0001; Partial vs Buried: *P*=0.0001). (**C**) There was no significant difference in egg hatching rates across all three categories of the remaining eggs after ant foraging (Surface vs. Partial: *P*=0.6613; Surface vs Buried: *P*=0.2665; Partial vs Buried: *P*=0.8782). (Kruskal-Wallis test with Dunn’s multiple comparison test). Bars represent mean ± S.E.M.

Given that laying eggs deeply into the substrate protects them from predation, why don’t fly mothers perform this behavior constitutively? We tested whether buried eggs suffer some fitness cost compared with eggs laid on the substrate surface. First, we tested whether there was any reduction in offspring developmental time or survival for flies that were buried in the egg stage, but did not find any difference in eclosion time or eclosion success between flies manually positioned on the food surface versus buried at the egg stage (Supp. Fig. 2A). We also tested whether the presence of older fly larvae churning the food, reduced temperature, or simulated rain (water misting) harmed the buried eggs, but once again found no difference in survival to eclosion (Supp. Fig. 2B-D). These results suggest that any cost of deep oviposition is not incurred by the offspring, but instead may be borne by fly mothers. The *D. melanogaster* oviposition program has been described as a progression of stereotyped phases starting with a searching phase, then the oviposition phase, and finally the clean-and-rest phase (Yang et al., 2008). We hypothesized that female flies might require more time to lay buried eggs or to rest after laying buried eggs. However, video analysis of the fly oviposition program during parasitoid wasp exposure showed no difference in the average length of either of these phases when the flies laid a surface egg versus laid a buried egg (Supp. Fig. 2E-F). Curiously, female flies spent significantly longer periods laying partially buried eggs as compared to surface eggs or fully buried eggs (Supp. Fig. 2E), suggesting that partially buried eggs need more care to position. In summary, we were unable to detect any negative consequences of the induced egg burying behavior. Either there is no fitness cost associated with this behavior, or there is a cost that we have not yet identified.

### Egg burying is a conserved trait across *Drosophila*

To test whether ant-induced egg burying is a conserved defensive behavior in flies, we assayed the behavior in three other *Drosophila* species of varying evolutionary distances from *D. melanogaster: D. simulans* (∼3MY since the most recent common ancestor), *D. yakuba* (∼5MY), and *D. virilis* (∼30MY) (Powell, 1997). Like *D. melanogaster*, the other *Drosophila* species maintained their egg production level when exposed to *F. mccooki* ants (Fig. 3A), and also like *D. melanogaster*, all three *Drosophila* species consistently laid their eggs deeper in the substrate in the presence of ants (Fig. 3B). This result holds even though *D. virilis*, and to some extent *D. yakuba*, tend to lay more deeply buried eggs than *D. melanogaster* in the unexposed state (Fig. 3B). These data suggest that although normal egg depth behavior varies across species, the switch to laying more buried eggs to prevent egg predation is a conserved trait.

**Fig. 3:**
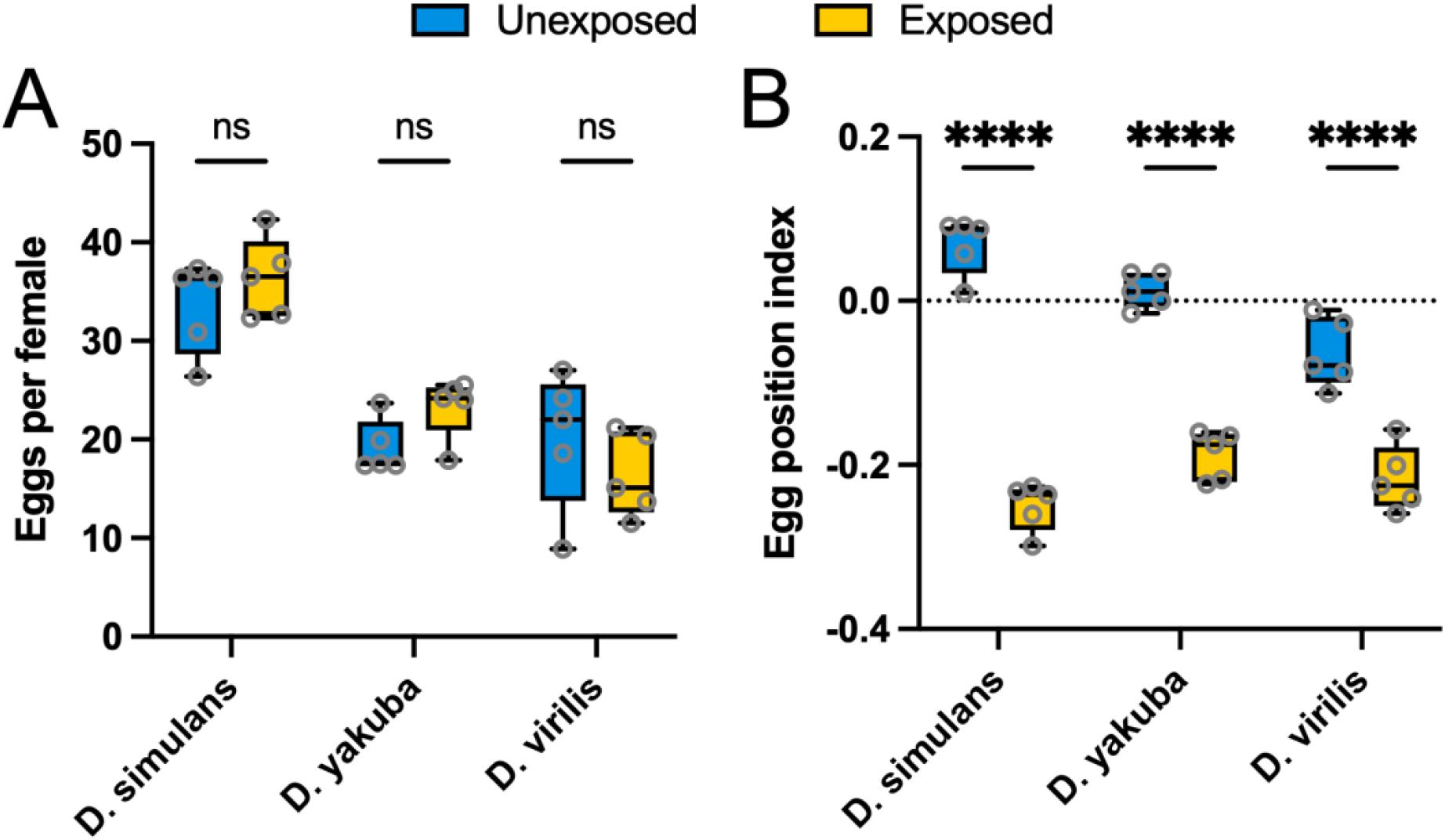
The induced egg burying behavior is conserved across Drosophilids. (**A**)Three other fly species did not reduce oviposition rates when exposed to *F. mccooki* ants (*D. simulans*: *P*=0.9729; *D. yakuba*: *P*=0.4829; *D. virilis*: *P*=0.6040). (**B**) All three species altered their oviposition depth when exposed to ants (*P*<0.0001 for all comparisons). (Two-way ANOVA with Bonferroni multiple comparison test).

### Olfaction mediates ant detection

Fly mothers sense and respond to the presence of ants by burying their eggs, but how do they know that ants are present? Flies have been shown to detect parasitoid wasps through the combined effects of the visual and olfactory systems (Ebrahim et al., 2015; Kacsoh et al., 2015, 2013; Lynch et al., 2016), and given that flies also bury their eggs in the presence of wasps (Supp. Fig. 1) we tested both of these sensory modalities. *GMR-hid* flies express a pro-apoptotic factor in the eye, making their eyes significantly reduced in size (Grether et al., 1995). These sight-deficient flies maintained their ability to respond to the presence of *F. mccooki* ants by burying their eggs (Fig. 4A). *GMR-hid* as well as *ninaB*^*1*^ mutant flies (which are blind due to loss of photoreceptors) also maintained their ability to sense and respond to *L. heterotoma* parasitoid wasps (Supp. Fig. 3A-B). These data suggest that vision is dispensable for the altered egg depth behavior. We next tested the necessity of the olfactory system: *orco*^*2*^ mutant flies lack the odorant receptor co-receptor and therefore lack most olfactory ability (Larsson et al., 2004). Heterozygous *orco*^*2*^ mutant flies showed a strong induced egg burying response in the presence of *F. mccooki* ants, but the anosmic homozygous *orco*^*2*^ mutant flies showed a much weaker (albeit still significant) response (Fig. 4B). Exposing the anosmic flies to either *P. hyatti* ants or *L. heterotoma* parasitoid wasps resulted in a complete failure to alter egg depth (Supp. Fig. 3C-D). To further test the necessity of olfaction for the egg burying behavior, we manually ablated the two main olfactory sensing organs from experimental flies either individually or in tandem. While ablating the maxillary palps had no effect on fly ability to bury eggs in the presence of *F. mccooki* ants, ablating the fly antennae abolished their ability to respond to the ants (Supp. Fig. 3E). Altogether, these data demonstrate that flies require olfactory input through their antennae to sense ants in their environment and alter oviposition depth behavior accordingly.

**Fig. 4:**
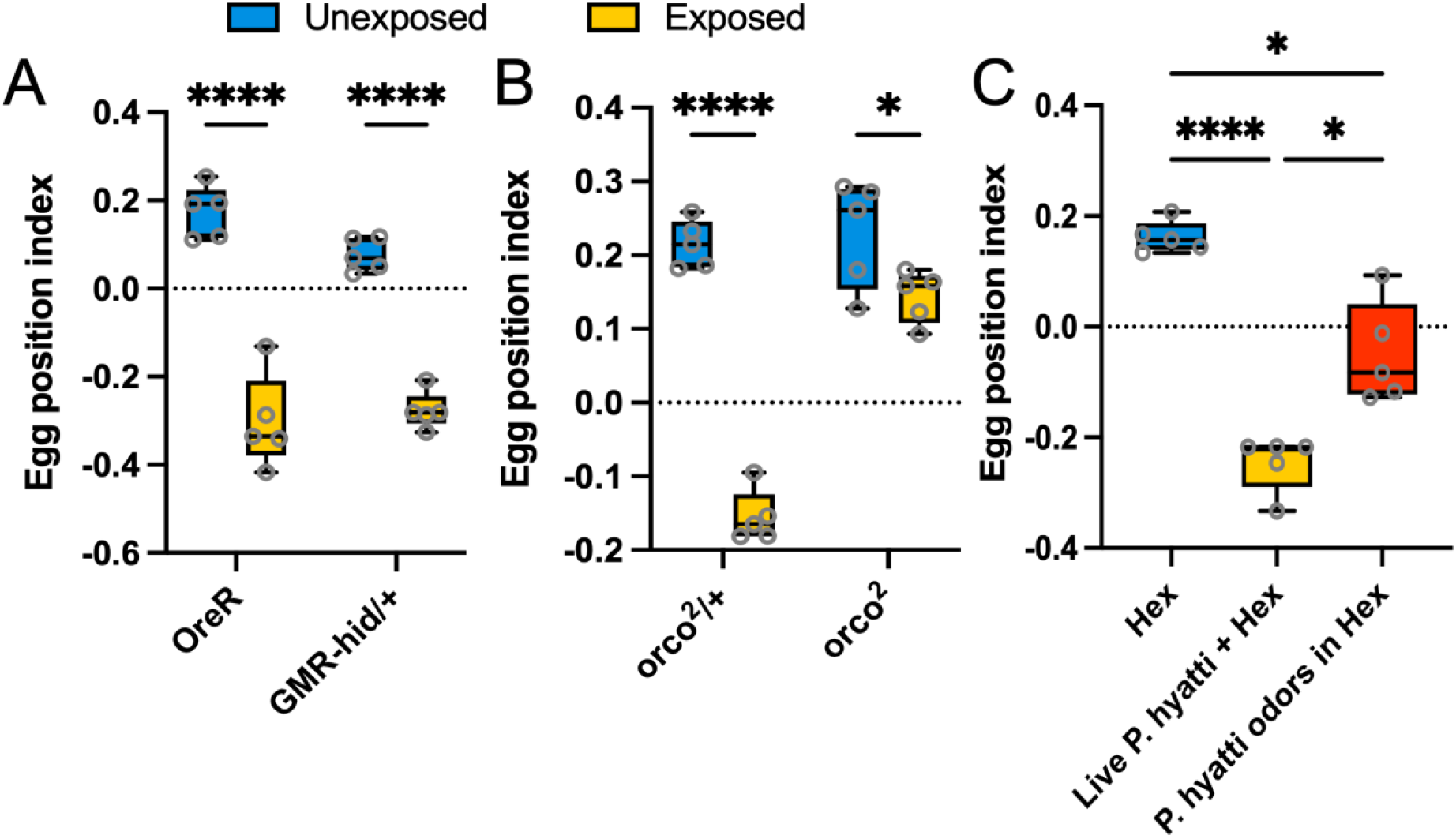
Flies detect ants through olfactory cues. (**A**) Blind flies continued to bury their eggs when exposed to *F. mccooki* ants, similar to control OreR flies (*P*<0.0001 for both comparisons). (**B**) Both control heterozygous *orco* mutant flies and homozygous *orco* mutants buried their eggs when exposed to ants, although the magnitude of the effect was much weaker for the homozygous mutants (*orco*^*2*^/+: *P*<0.0001; *orco*^*2*^: *P*=0.0210; two-way ANOVA with Bonferroni multiple comparison test for A and B). (**C**) Ant (*P. hyatt*i) odors extracted in hexane significantly altered fly oviposition depth behavior, although not to the same extent as direct ant exposure (Hex vs live ants: *P*<0.0001; Hex vs ant odors: *P*=0.0166; Live ants vs ant odors: *P*=0.0150; Brown-Forsythe and Welch’s ANOVA with Dunnett’s multiple comparison test).

It was previously shown that the fly odorant receptors Or49a and Or85f, which are expressed in the same olfactory receptor neurons in adult flies, detect specific odorants from parasitoid wasps in the genus *Leptopilina*, in turn driving avoidance behaviors during fly oviposition (Ebrahim et al., 2015). One of these compounds, iridomyrmecin, was first isolated from the Argentine Ant, *Linepithema humile* (formerly *Iridomyrmex humilis*) (Pavan, 1948) suggesting flies may also detect ants via this mechanism. Using an *Or49a-Gal4* driver, we inhibited activity of Or49a/Or85a-expressing olfactory receptor neurons by driving expression of *Kir2*.*1* (which causes neuronal membrane hyperpolarization) in these neurons, or we completely ablated these neurons by expressing in them the proapoptotic gene *hid*. Neither modification blocked the fly egg burying behavior in the presence of *L. heterotoma* parasitoid wasps (Supp. Fig. 3F-G). These data suggest that flies are using a different type of olfactory sensory neuron and olfactory receptor to detect a novel odorant associated with ants.

If an ant olfactory stimulus is sufficient to induce the fly oviposition depth switch, we hypothesized that ant body-wash extracts could replace live ants as the egg burying stimulus. We used the solvent hexane to extract odorants from the bodies of *P. hyatti* ants and added the odors directly onto the food substrate. While flies that were exposed to live *P. hyatti* ants as a positive control showed a strong induced egg burying behavior, flies exposed to ant odors alone showed a weaker, though still significant, response (Fig. 4C). There are at least three reasons why odor extracts do not fully recapitulate the direct ant exposure condition. First, it is possible that the body wash extracts only possess a subset of multiple distinct odorants required for the full behavioral response. To test this, we used different solvents that varied in their polarity to target different molecules. Interestingly, extracts collected in non-polar hexane (Fig 4C, Supp. Fig. 4A) and dichloromethane (Supp.Fig. 4B) were both sufficient to induce the partial egg burying behavior, while use of more strongly polar solvents failed to induce any fly response (Supp. Fig. 4C-E). However, combining body wash odorants extracted using hexane and dichloromethane did not induce the full egg burying behavior (data not shown), showing that no additive effect is achieved by combining the odorant subsets extracted by each solvent. Second, it is possible that exposure to insect body wash extracts fails to fully recapitulate the live insect exposure results because the body wash odorants dissipate over the course of the experiment without being replaced. To test this idea, we exposed flies to both dead *L. heterotoma* parasitoid wasps, and to vials in which the wasps had been housed, both treatments of which should have the full wasp odorant profile but no new odorants being produced. In both cases, flies showed a partial egg burying response similar to their response to insect body wash extracts, supporting the odorant dissipation hypothesis (data not shown). Third, it is possible that flies are using some other sensory modality besides olfaction or vision to detect ant/wasp presence, such as gustation, a hypothesis that remains untested. Regardless, we were able to extract an odorant from insect bodies that flies detected, causing them to induce at least a partial egg burying response.

### Flies respond to most hymenopterans

We have shown that flies lay eggs more deeply into the food substrate when exposed to ants (Fig. 1), and that this behavior protects the eggs from ant predation (Fig. 2). Flies also induce this behavior when exposed to a parasitoid wasp, although it is unclear what benefit there is to burying eggs in this context given that *L. heterotoma* does not attack fly offspring until they reach the larval stage, when they are crawling on top of the food. To determine the breadth of ants and potentially other insects that induce flies to bury their eggs, we exposed flies to body washes or live samples of numerous arthropods. Flies responded to the vast majority of ant species tested representing the three ant subfamilies Dolichoderinae, Formicinae, and Myrmicinae (Fig. 5). Exceptions included the Myrmicinae species *Monomorium cyaneum*, as well as some trials from several other ant species (Fig. 5). Parasitoid wasps represent another large branch of hymenopterans that impose strong selective pressures on fly offspring survival. Flies that were exposed directly to parasitoid wasps that infect fly larvae (genus *Leptopilina, Ganaspis*, and *Asobara*) or fly pupae (*Pachycrepoideus* and *Trichopria*) significantly altered their oviposition depth (Fig. 5). However, flies also responded to the body wash extract of the whitefly parasitoid, *Encarsia inaron*, as well as extracts from every other hymenopteran tested (bees and paper wasps), even though *D. melanogaster* has no known ecological interactions with these insects (Fig. 5).

**Fig. 5:**
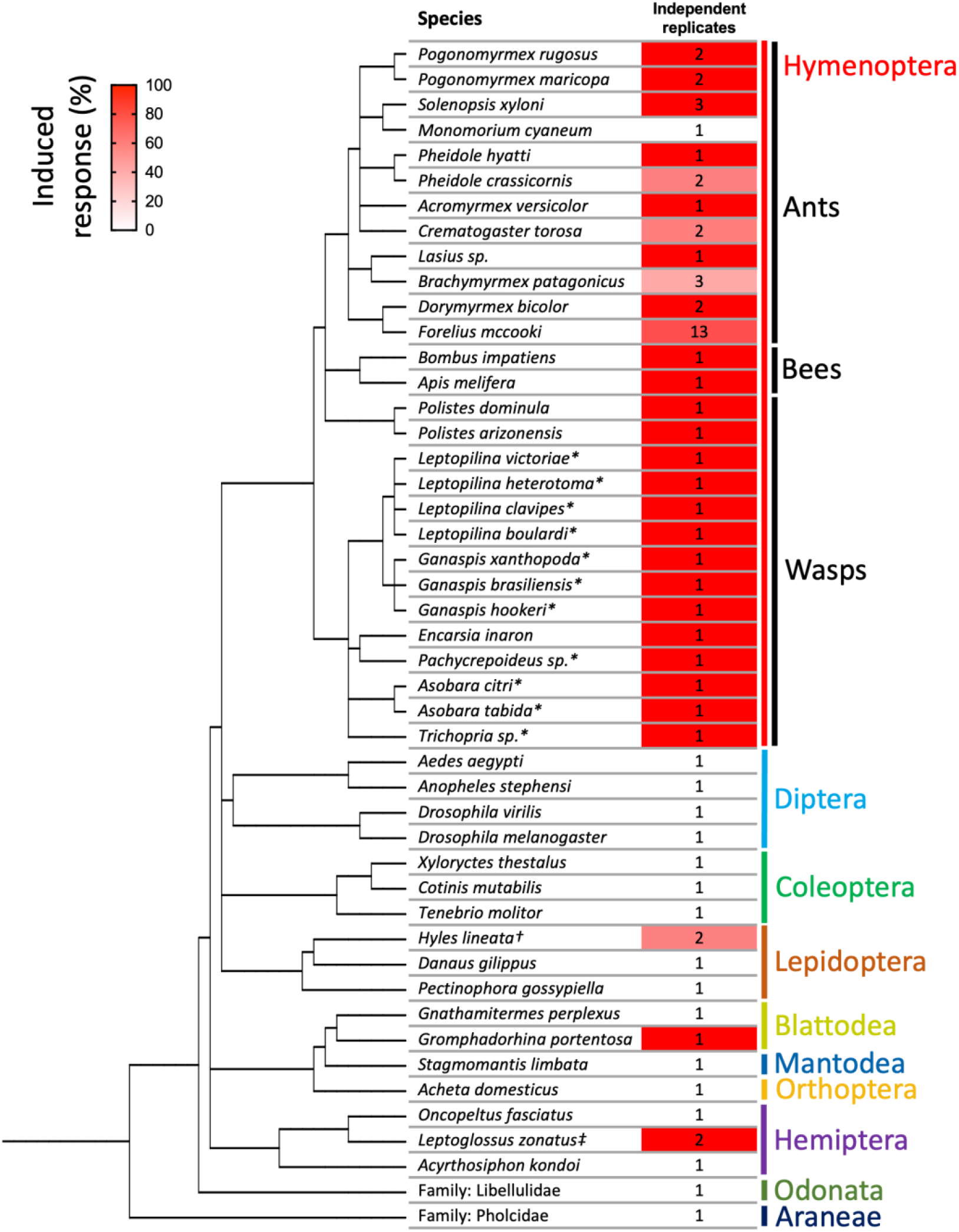
Hymenopteran odors trigger the fly egg burying behavior. A phylogeny is shown of the different insect species whose odors were tested for the ability to induce the fly egg burying behavior. The number of independent replicates run for each insect species is shown, while the percentage of replicates resulting in significant fly egg burying is colored in red. Flies responded to odors from a diversity of hymenopterans, but rarely to odors from non-hymenopterans. * live insect exposure rather than body wash exposures; † odor extracted from larval or adult stage; ‡ unlike the ant replicates that were derived from independent ant colonies, the *Leptoglossus zonatus* replicates were derived from the same colony, although the odor extractions were independent.

The odor extracts from non-hymenopteran insects generally did not induce flies to alter their oviposition behavior, but there were three exceptions. First, odor extract from the leaf-footed bug, *Leptoglossus zonatus*, did induce the fly egg burying behavior in two separate trials, despite body washes from other hemipterans showing no effect on the flies (Fig. 5). Second, odor extract from the nymph stage of the Madagascar hissing cockroach, *Gromphadorhina portentosa*, also affected fly oviposition behavior. Third, odors from the adult stage of the moth *Hyles lineata* altered fly oviposition behavior, but odors from the larval stage had no such effect. In sum, flies responded to odors from 27 of 28 hymenopteran insects, but only 3 of 19 non-hymenopteran insects. Our data suggest that an odorant evolved early in the hymenopterans and has been maintained over time, with sporadic gains and losses in the broader insect group, and that flies evolved to sense this odorant as a means of protecting their offspring from predatory ants.

## Discussion

Recognition of environmental threats, and execution of an appropriate response, is critical to organismal survival. Here, we have shown that flies detect ant presence through olfactory sensing of hymenopteran odors, and switch their oviposition behavior to more deeply position their eggs in the food substrate. This protects fly eggs from ant predation.

Egg laying behavior follows a similar stereotyped pattern across different fly species. It is first characterized by a search-like, informational gathering processes about the local nutrient availability and substrate stiffness through labellum and leg contact (Bräcker et al., 2019; Yang et al., 2008; Zhang et al., 2020). Once a suitable site is found, flies switch to a more refined search behavior, including ovipositor contact with the substrate (Bräcker et al., 2019), before an egg is finally laid. These discrete micro-behaviors are conserved across *Drosophila*, with minor variations depending on the ecology of each species. For example, the agricultural pest *D. suzukii* spends significantly more time than *D. melanogaster* contacting the food substrate with its ovipositor and expelling its eggs (Bräcker et al., 2019), presumably because it tends to lay eggs more deeply into less ripened fruits (Karageorgi et al., 2017). Furthermore, *Drosophila* species that oviposit on mushrooms tend to deeply bury their eggs (Rouquette & Davis, 2003), perhaps because mushroom flesh is soft, or because ants commonly visit mushrooms (Lewis & Worthen, 1992). While fly oviposition behaviors are well-studied (Bräcker et al., 2019; Cury et al., 2019; Karageorgi et al., 2017; Van Mele et al., 2009; Wang et al., 2020; Yang et al., 2008; Zhang et al., 2020), this is the first report describing an induced change in oviposition depth.

Our measure of oviposition depth, the egg position index, is a blunt tool. While egg depth is a continuous trait, we bin the eggs into one of three depth categories, and most of the eggs are classified as ‘partially buried’. A lot of the behavior may be missed when an egg that is only 20% inserted into the substrate is scored the same as an egg that is 80% inserted, especially if these eggs have different levels of protection against foraging ants. Ideally a more sensitive measure of egg depth could be devised. Nevertheless, the fact that we see significant differences in the egg position index across treatments indicates that egg burying is a robust and important fly behavior.

Several examples exist of organisms altering oviposition behavior in response to biotic threats. For example, water striders (*Aquarius paludum insularis*) oviposit their eggs deeper in the water column after exposure to parasitoid wasps, which limits egg parasitism (Hirayama & Kasuya, 2009). Newts (*Taricha granulosa*) lay their eggs attached to plants higher in the water column to avoid egg predator caddisfly larvae (*Limnephilus flavastellus*) (Gall et al., 2012). There are even examples in fruit flies: *Drosophila* species avoid ovipositing at sites infested by toxic microbes, which they detect via the odorant geosmin (Stensmyr et al., 2012). They also alter oviposition in the presence of parasitoid wasps (Carton et al., 1986) using both visual and olfactory cues: female flies lay fewer eggs during forced exposures, they preferentially choose non-infested sites when given a choice by sensing the wasp odorant iridomyrmecin, and they preferentially lay their eggs in more toxic (alcoholic) environments when exposed (Kacsoh et al., 2013; Lefèvre et al., 2012; Lynch et al., 2016; Ebrahim et al., 2015). All of these are examples of threat-induced behavioral changes that organisms make about ‘when’ or ‘where’ to oviposit, whereas the new behavior described here is about ‘how’ flies choose to oviposit. Furthermore, we know that the fly responses to geosmin and iridomyrmecin are hardwired into the fly, using specific olfactory receptors and neuronal circuits in the brain. It will be interesting to determine how these different environmental cues are signaled through the brain to regulate different types of egg laying behaviors, and whether there is any overlap in each circuit.

Ants are a ubiquitous presence across diverse habitats, and many insects have evolved ant avoidance behaviors triggered by distinct sensory modalities. For example, honeybees avoid flowers in the field that have live ants or their odors nearby (Li et al., 2014; Sidhu & Rankin, 2016), and tephritid fruit flies avoid fruits perfumed with ant pheromones (Van Mele et al., 2009). Furthermore, butterflies (*Eunica bechina*) were reported to avoid oviposition sites due to visual detection of ant presence, although the fact that ants were pinned to plant leaves suggests that ant odors may also have contributed to the butterfly avoidance (Sendoya et al., 2009). Surprisingly, termites (*Coptotermes acinaciformis*) were shown to detect and avoid ants based on the vibrational pattern of ants walking through their woody substrates (Oberst et al., 2017). This may reflect a keen sense of vibration detection in termites, which have been shown to communicate alarm signals via similar vibrational cues (Inta et al., 2009). These examples show that detection of ant threats is common in the insect world, and that insects have evolved diverse sensory mechanisms for identifying ant presence. Given that *D. melanogaster* invests more resources into its olfactory system than other sensory systems (Keesey et al., 2019), it may not be surprising that they detect ant predators through olfactory signals. In the future, it will be interesting to identify the specific hymenopertan odorant and the fly odorant receptor(s) responsible for sensing of ant odors and the neural circuitry that responds to odor sensing by altering oviposition choices (Wang et al., 2020).

Although olfaction is necessary for fly detection of ants, we could not prove that ant odors are sufficient to induce the full fly egg burying phenotype (Fig. 4). We believe there are two likely reasons why providing flies with ant body wash odorant extracts does not fully recapitulate the direct insect-exposure results. First, the ant odor extract experiments likely started with a high concentration of ant odors, but over the course of the 24 hour experiment these volatile odorant concentrations may have declined to a point where fly behavior was no longer altered. In contrast, the live-ant exposures may have maintained a high concentration of odorants throughout the experiments due to constant release from the insects. Second, it is possible that fly oviposition behaviors like egg burying require multimodal sensory integration such that ant odors suppress the preference for ‘surface’ eggs, but a second stimulus enhances the magnitude of preference for ‘buried’ eggs. This second stimulus might include gustatory or auditory cues from the ants. This kind of gating mechanism has been observed for fly larval rolling behavior during parasitoid attack, where vibrational cues enhance the fly rolling response to mechanical poking by the wasp ovipositor (Ohyama et al., 2015). Further testing of fly sensory systems will help identify additional inputs into the oviposition depth behavioral response.

We showed that the ability of flies to sense ant odorants, and induce the egg burying behavior, is beneficial to the flies due to the increased survival of their eggs during ant predation events (Fig. 2). However, flies also induce the egg burying behavior in response to the odors of other hymenopterans like wasps and bees, even though wasps and bees are not known to harm fly eggs. This suggests that some conserved odorant dates back to the common ancestor of all hymenopterans. Likewise, the ant odor-induced egg burying behavior is conserved across the genus Drosophila (Fig. 3). Given that the expansion of ant, wasp, and bee lineages seems to coincide with the expansion of dipterans in the Jurassic period over 150 million years ago (Misof et al., 2014; Peters et al., 2017), it is possible that fly recognition of hymenopteran threats is ancient. In the future, it will be interesting to determine if other dipterans mount behavioral defenses in response hymenopteran threats, and if the neurogenetic basis of this sensing is conserved across dipterans.

## Acknowledgments

We would like to thank Goggy Davidowitz, Anna Dornhaus, Martha Hunter, Michael Riehle, Patricia Stock, Bruce Tabashnik, and Kathleen Walker from the University of Arizona, Vanessa Corby-Harris from the USDA, and Elizabeth Tibbetts from the University of Michigan for providing insects for odor extraction. This research was funded by the University of Arizona BIO5 Institute Postdoctoral Fellowship to SMD and NSF grant IOS-1257469 to TAS. The authors acknowledge resources and support from the Knowledge Enterprise Biosciences Mass Spectrometry Core Facility at Arizona State University.

---

**Supp.Fig. 1:**
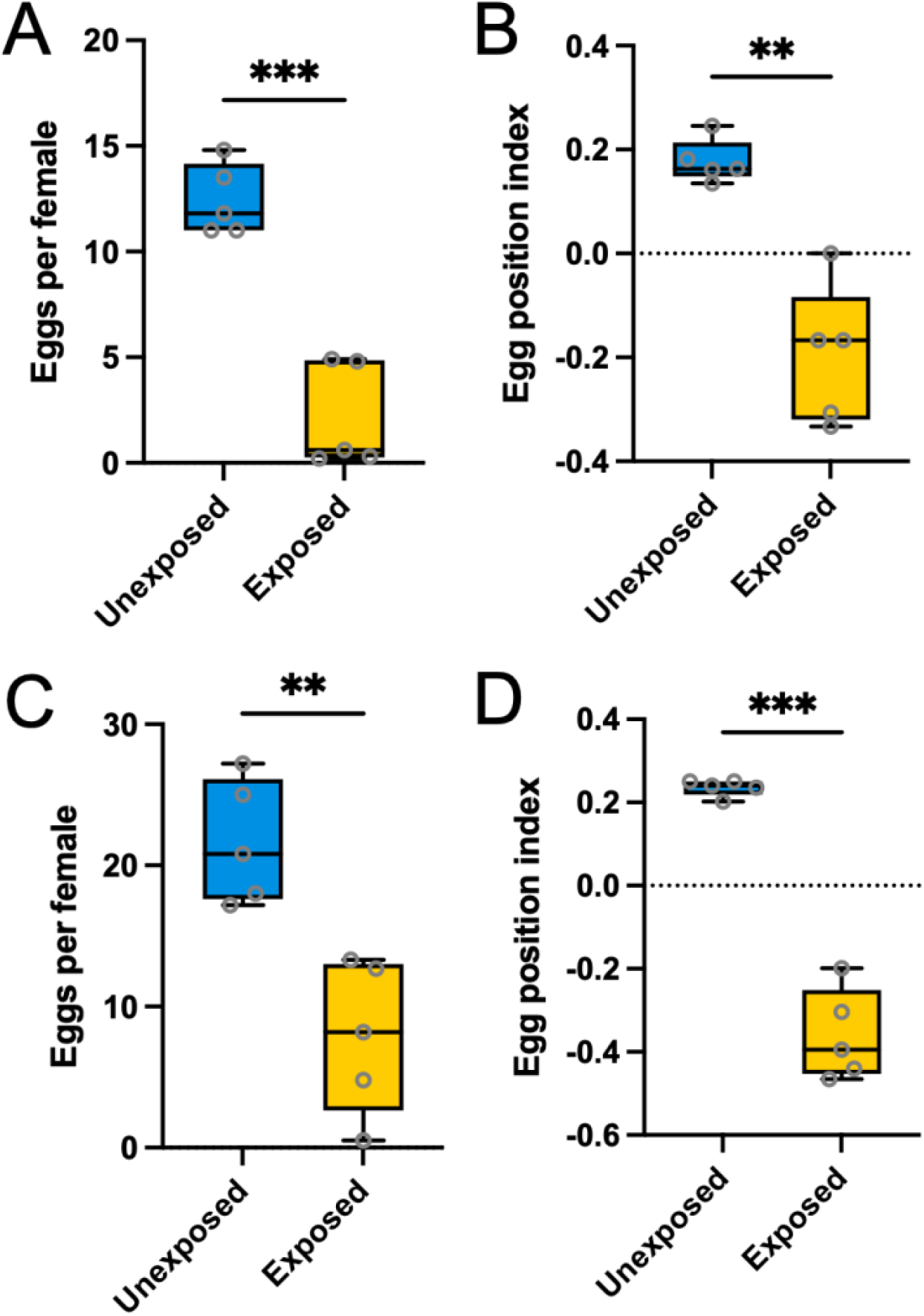
Exposure to parasitoid wasps also alters fly oviposition behaviors. Unlike their response to ant exposure, flies reduced oviposition rates when exposed to *L. heterotoma* (**A**, *P*=0.0001) or *L. boulardi* (**C**, *P*=0.0025) parasitoid wasps. However, like their response to ant exposure, flies oviposited eggs more deeply when exposed to *L. heterotoma* (**B**, *P*=0.0022) and *L. boulardi* (**D**, *P*=0.0002) wasps. (Unpaired, Welch’s t-test).

**Supp.Fig. 2:**
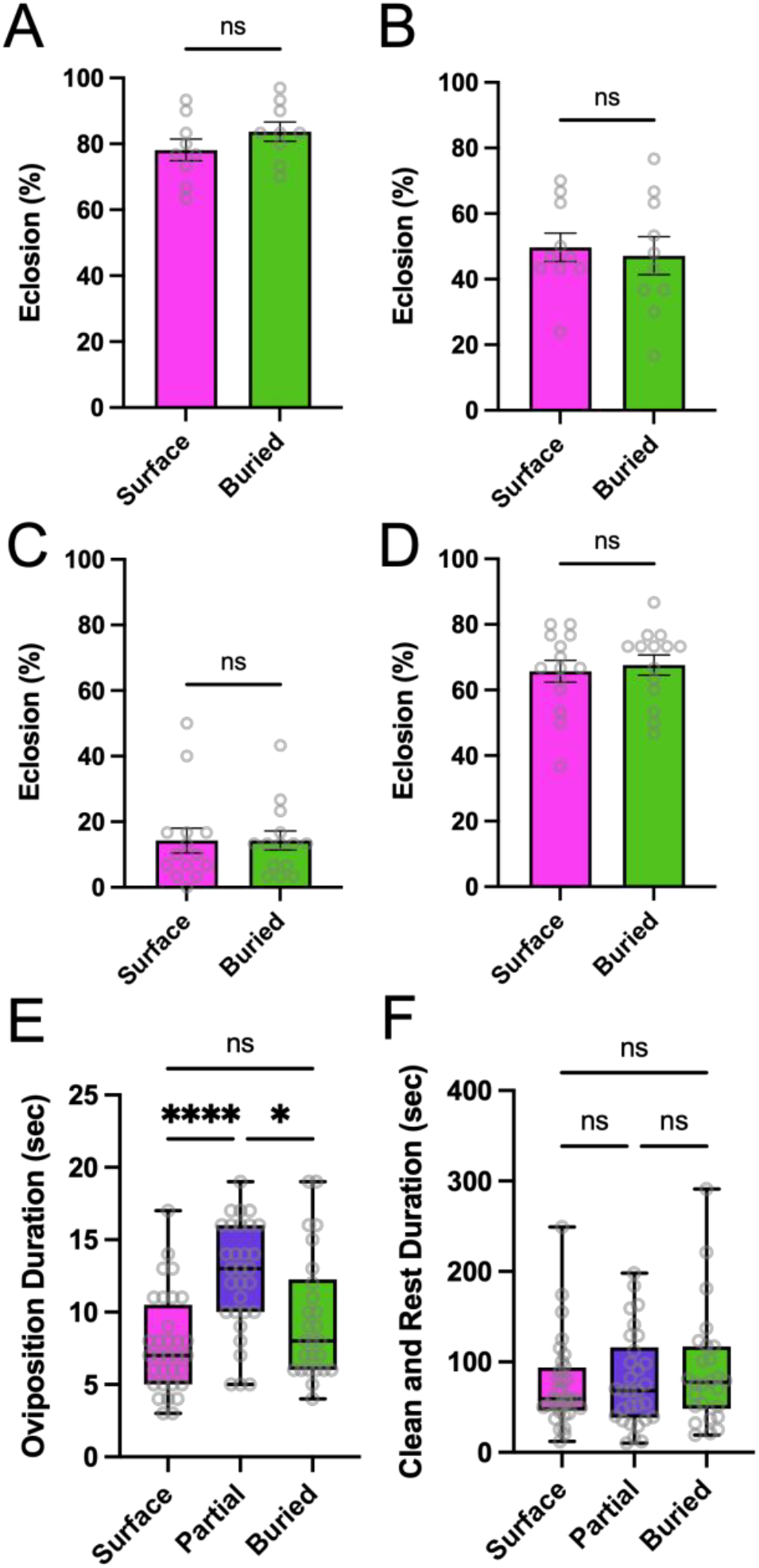
Any cost to deeper oviposition remains unknown. Egg depth had no effect on adult eclosion rates under normal conditions (**A**, *P*=0.2230) or when other survival risks were present, such as the presence of older larvae (**B**, *P*=0.7234), cold overnight temperatures (**C**, *P*=0.7577), or simulated rain (**D**, *P*=0.6774) (unpaired, Welch’s t-test for A, B, and D; Mann-Whitney test for C). (**E**) Maternal oviposition duration was significantly increased for partially inserted eggs, but no timing difference was found between surface and buried eggs (Surface vs. Partial: *P*<0.0001; Surface vs Buried: *P*<0.3475; Partial vs Buried: *P*=0.0386). (**F**) After ovipositing, the female flies showed no difference in the length of their clean-and-rest phase across the three egg position categories (Surface vs. Partial: *P*=0.; Surface vs Buried: *P*<0.; Partial vs Buried: *P*=0.). (Kruskal-Wallis with Dunn’s multiple comparison test in E and F). Bars represent mean ± S.E.M.

**Supp.Fig. 3:**
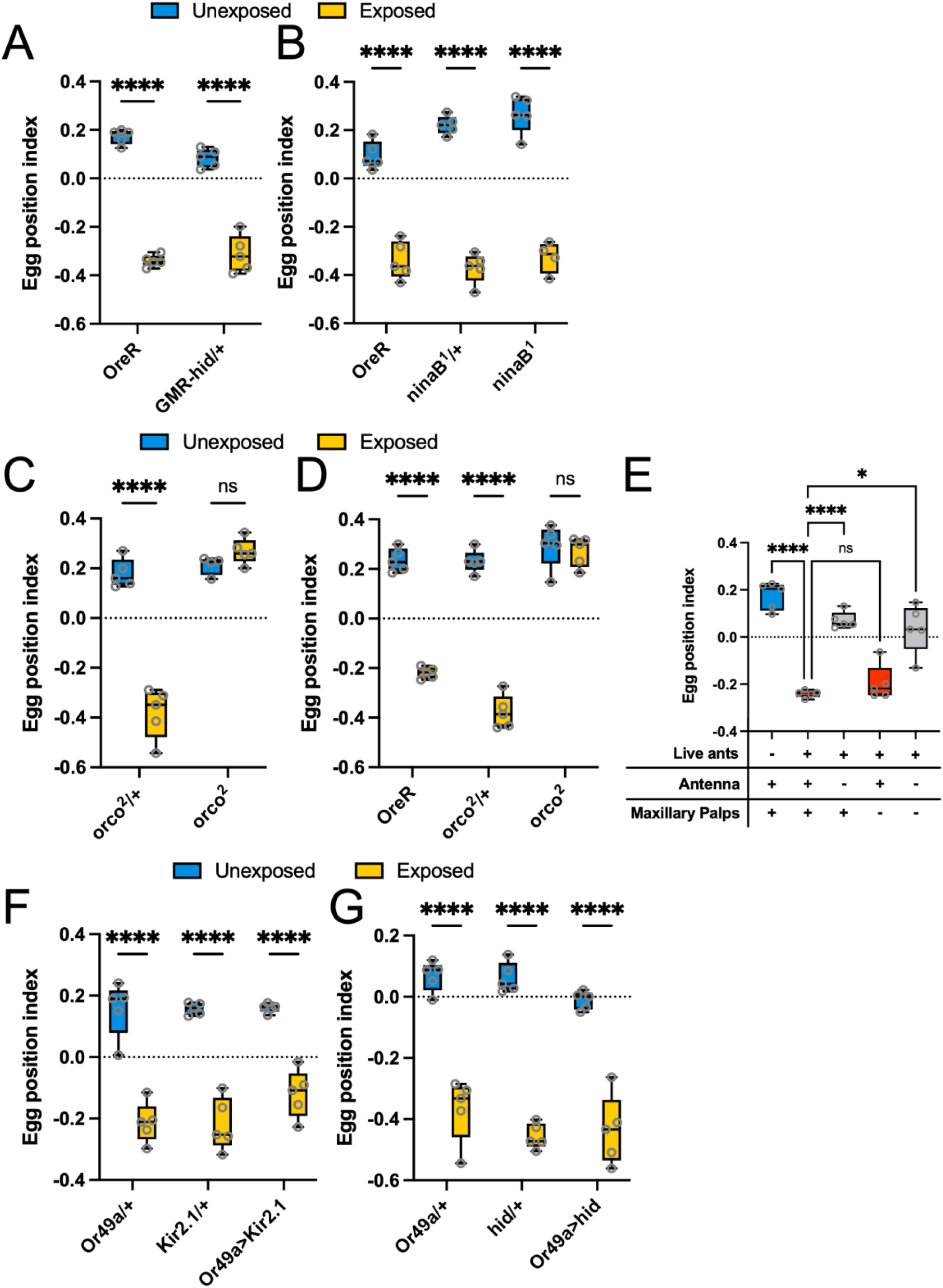
Olfaction is required to detect ants and wasps. (**A** and **B**) Blind flies exposed to *L. heterotoma* parasitoid wasps maintained normal oviposition depth change similar to control OreR flies or heterozygous controls (*P*<0.0001 for all comparisons in A and B). (**C**) Anosmic *orco* mutant flies failed to alter oviposition depth behavior when exposed to *P. hyatti* ants (*orco*^*2*^/+: *P*<0.0001; *orco*^*2*^: *P*=0.4805). (**D**) Similarly, anosmic flies also failed to respond to *L. heterotoma* wasps (OreR: *P*<0.0001; *orco*^*2*^/+: *P*<0.0001; *orco*^*2*^: *P*>0.9999). (**E**) Control OreR flies with ablated antenna had a significantly higher egg position index when exposed to *F. mccooki* ants than fully intact flies. Ant-exposed flies with only maxillary palp ablations showed no significant difference in oviposition depth compared to intact flies (intact exposed vs intact unexposed: *P*<0.0001; intact exposed vs antenna-ablated exposed: *P*<0.0001; intact exposed vs maxillary palp-ablated exposed: *P*=0.6090; intact exposed vs both-ablated exposed: *P*=0.0138). (**F** and **G**) Flies responded to *L. heterotoma* wasp presence by burying eggs even with the silencing (*Kir2*.*1*) or ablation (*hid*) of wasp odor-detecting Or49a sensory neurons (*P*<0.0001 for all comparisons in F and G). (A-D, F, G: Two-way ANOVA with Bonferroni multiple comparison test; E: Brown-Forsythe and Welch’s ANOVA with Dunnett’s multiple comparison test).

**Supp.Fig. 4:**
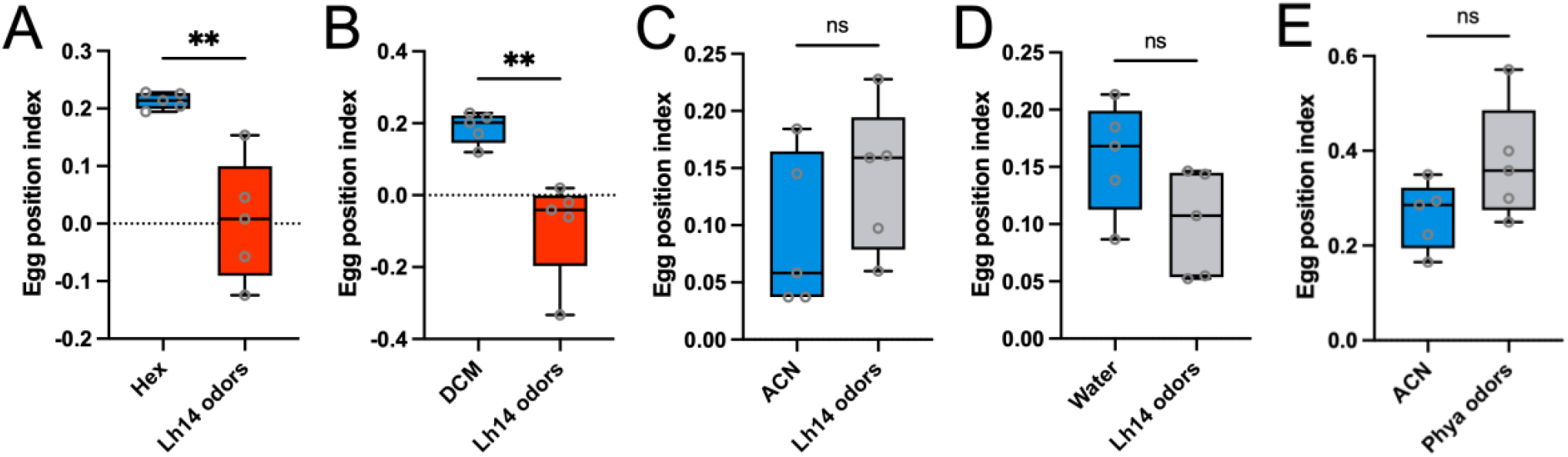
Body wash solvent specificity in fly detection of ant and wasp odors. Different solvents used to extract body odors varied in their success at inducing the fly egg burying behavior. Flies responded to *L. heterotoma* parasitoid wasp body washes when either hexane (Hex) (**A**, *P*=0.0110) or dichloromethane (DCM) (**B**, *P*=0.0099) were used. Body washes using other solvents, such as acetonitrile (ACN) (**C**, *P*=0.2802) and water (**D**, *P*=0.0899), did not induce the fly behavior. *P. hyatti* ant body washes using the ACN solvent also showed no significant change in oviposition behavior (**E**, *P*=0.1249). (Unpaired, Welch’s t-test).

## References

Ache, J. M., Polsky, J., Alghailani, S., Parekh, R., Breads, P., Peek, M. Y., Bock, D. D., Von Reyn, C. R., & Card, G. M. (2019). Neural Basis for Looming Size and Velocity Encoding in the Drosophila Giant Fiber Escape Pathway. Current Biology, 29, 1073–1081. https://doi.org/10.1016/j.cub.2019.01.079

Baker, B. S., Taylor, B. J., & Hall, J. C. (2001). Are Complex Behaviors Specified by Dedicated Regulatory Genes? Reasoning from Drosophila. Cell, 105(1), 13–24. https://doi.org/10.1016/S0092-8674(01)00293-8

Borne, F., Prigent, S. R., Molet, M., & Courtier-Orgogozo, V. (2021). Drosophila glue protects from predation. Proc. R. Soc. B, 288: https://doi.org/10.1098/rspb.2021.0088

Bowman, W. D., & Hacker, S. D. (2020). Predation. In Ecology (5th ed.). Oxford University Press.

Bräcker, L. B., Schmid, C. A., Bolini, V. A., Holz, C. A., Prud’homme, B., Sirota, A., & Gompel, N. (2019). Quantitative and discrete evolutionary changes in the egg-laying behavior of single Drosophila females. Frontiers in Behavioral Neuroscience, 13. https://doi.org/10.3389/fnbeh.2019.00118

Carton, Y., Bouletreau, M., Alphen, J. J. M. van, & Lenteren, J. C. van. (1986). The Drosophila parasitic wasps. In M. Ashburner, E. Novitski, H. L. Carson, & J. N. Thompson (Eds.), The Genetics and Biology of Drosophila (pp. 347–394). Academic Press.

Chen, J. (2017). Freeze–Thaw Sample Preparation Method Improves Detection of Volatile Compounds in Insects Using Headspace Solid-Phase Microextraction. Analytical Chemistry, 89(16), 8366–8371. https://doi.org/10.1021/ACS.ANALCHEM.7B01622

Chowdhury, B., Wang, M., Gnerer, J. P., & Dierick, H. A. (2021). The Divider Assay is a high-throughput pipeline for aggression analysis in Drosophila. Communications Biology, 4(1). https://doi.org/10.1038/S42003-020-01617-6

Combes, S. A., Rundle, D. E., Iwasaki, J. M., & Crall, J. D. (2012). Linking biomechanics and ecology through predator–prey interactions: flight performance of dragonflies and their prey. Journal of Experimental Biology, 215(6), 903–913. https://doi.org/10.1242/JEB.059394

Cury, K. M., Prud’homme, B., & Gompel, N. (2019). A short guide to insect oviposition: when, where and how to lay an egg. Journal of Neurogenetics, 33(2), 75–89. https://doi.org/10.1080/01677063.2019.1586898

de la Flor, M., Chen, L., Manson-Bishop, C., Chu, T. C., Zamora, K., Robbins, D., Gunaratne, G., & Roman, G. (2017). Drosophila increase exploration after visually detecting predators. PloS ONE, 12(7), 1–17. https://doi.org/10.1371/journal.pone.0180749

Ebrahim, S. A., Dweck, H. K., Stokl, J., Hofferberth, J. E., Trona, F., Weniger, K., Rybak, J., Seki, Y., Stensmyr, M. C., Sachse, S., Hansson, B. S., & Knaden, M. (2015). Drosophila avoids parasitoids by sensing their semiochemicals via a dedicated olfactory circuit. PLOS Biology, 13(12), e1002318. https://doi.org/10.1371/journal.pbio.1002318

Escalante, A., & Benado, M. (1990). Predation on the Cactophilic Fly, Drosophila starmeri, in the Columnar Cactus, Pilosocereus lanuginosus. Biotropica, 22(1), 48–50.

Fernandes, W. D., Sant’Ana, M. V., Raizer, J., & Lange, D. (2012). Predation of fruit fly larvae Anastrepha (Diptera: Tephritidae) by ants in grove. Psyche, 2012, 1–7. https://doi.org/10.1155/2012/108389

Fisher, B. L., & Cover, S. P. (2007). Ants of North America : a Guide to the Genera. University of California Press.

Folmer, O., Black, M., Hoeh, W., Lutz, R., & Vrijenhoek, R. (1994). DNA primers for amplification of mitochondrial cytochrome c oxidase subunit I from diverse metazoan invertebrates. Molecular Marine Biology and Biotechnology, 3(5), 294–299.

Freitas, A. V. L., & Oliveira, P. S. (1996). Ants as Selective Agents on Herbivore Biology: Effects on the Behaviour of a Non-Myrmecophilous Butterfly. Journal of Animal Ecology, 65(2), 205–210. https://doi.org/10.2307/5723

Gall, B. G., Brodie III, E. D., & Brodie Jr, E. D. (2012). Fine-scale selection by ovipositing females increases egg survival. Ecology and Evolution, 2(11), 2763–2774. https://doi.org/10.1002/ece3.389

Grether, M. E., Abrams, J. M., Agapite, J., White, K., & Steller, H. (1995). The head involution defective gene of Drosophila melanogaster functions in programmed cell death. Genes and Development, 9(14), 1694–1708. https://doi.org/10.1101/gad.9.14.1694

Hermann, S. L., & Landis, D. A. (2017). Scaling up our understanding of non-consumptive effects in insect systems. Current Opinion in Insect Science, 20, 54–60. https://doi.org/10.1016/j.cois.2017.03.010

Hirayama, H., & Kasuya, E. (2009). Oviposition depth in response to egg parasitism in the water strider: high-risk experience promotes deeper oviposition. Animal Behaviour, 78(4), 935–941. https://doi.org/10.1016/j.anbehav.2009.07.019

Hölldobler, B., & Wilson, E. O. (1990). The ants. Belknap Press of Harvard University Press.

Inta, R., Evans, T. A., & Lai, J. C. S. (2009). Effect of vibratory soldier alarm signals on the foraging behavior of subterranean termites (Isoptera: Rhinotermitidae). Journal of Economic Entomology, 102(1), 121–126. https://doi.org/10.1603/029.102.0117

Kacsoh, B. Z., Bozler, J., Ramaswami, M., & Bosco, G. (2015). Social communication of predator-induced changes in Drosophila behavior and germline physiology. ELife, 4, 1–36. https://doi.org/10.7554/eLife.07423

Kacsoh, B. Z., Lynch, Z. R., Mortimer, N. T., & Schlenke, T. A. (2013). Fruit flies medicate offspring after seeing parasites. Science, 339(6122), 947–950. https://doi.org/10.1126/science.1229625

Karageorgi, M., Bräcker, L. B., Lebreton, S., Minervino, C., Cavey, M., Siju, K. P., Grunwald Kadow, I. C., Gompel, N., & Prud’homme, B. (2017). Evolution of Multiple Sensory Systems Drives Novel Egg-Laying Behavior in the Fruit Pest Drosophila suzukii. Current Biology, 27(6), 847–853. https://doi.org/10.1016/J.CUB.2017.01.055

Keesey, I. W., Grabe, V., Gruber, L., Koerte, S., Obiero, G. F., Bolton, G., Khallaf, M. A., Kunert, G., Lavista-Llanos, S., Valenzano, D. R., Rybak, J., Barrett, B. A., Knaden, M., & Hansson, B. S. (2019). Inverse resource allocation between vision and olfaction across the genus Drosophila. Nature Communications 2019 10:1, 10(1), 1–16. https://doi.org/10.1038/s41467-019-09087-z

Larsson, M. C., Domingos, A. I., Jones, W. D., Chiappe, M. E., Amrein, H., & Vosshall, L. B. (2004). Or83b encodes a broadly expressed odorant receptor essential for Drosophila olfaction. Neuron, 43(5), 703–714. https://doi.org/10.1016/j.neuron.2004.08.019

Lefèvre, T., de Roode, J. C., Kacsoh, B. Z., & Schlenke, T. A. (2012). Defence strategies against a parasitoid wasp in Drosophila: Fight or flight? Biology Letters, 8(2), 230–233. https://doi.org/10.1098/rsbl.2011.0725

Lemaitre, B., & Hoffmann, J. (2007). The Host Defense of Drosophila melanogaster. Annual Review of Immunology, 25, 697–743. https://doi.org/10.1146/ANNUREV.IMMUNOL.25.022106.141615

Letunic, I., & Bork, P. (2021). Interactive Tree Of Life (iTOL) v5: an online tool for phylogenetic tree display and annotation. Nucleic Acids Research, 49(W1), W293–W296. https://doi.org/10.1093/NAR/GKAB301

Lewis, G. P., & Worthen, W. B. (1992). Effects of Ant Predation and Mushroom Desiccation on the Survival of Mycophagous Drosophila tripunctata Larvae. Oikos, 64(3), 553–559. Retrieved from https://www.jstor.org/stable/3545175

Li, J., Wang, Z., Tan, K., Qu, Y., & Nieh, J. C. (2014). Giant Asian honeybees use olfactory eavesdropping to detect and avoid ant predators. Animal Behaviour, 97, 69–76. https://doi.org/10.1016/j.anbehav.2014.08.015

Lynch, Z. R., Schlenke, T. A., & de Roode, J. C. (2016). Evolution of behavioural and cellular defences against parasitoid wasps in the Drosophila melanogaster subgroup. Journal of Evolutionary Biology, 29(5), 1016–1029. https://doi.org/10.1111/jeb.12842

Markow, T. A. (2015). The secret lives of Drosophila flies. ELife, 4(JUNE). https://doi.org/10.7554/ELIFE.06793

Misof, B., Liu, S., Meusemann, K., Peters, R. S., Donath, A., Mayer, C., Frandsen, P. B., Ware, J., Flouri, T., Beutel, R. G., Niehuis, O., Petersen, M., Izquierdo-Carrasco, F., Wappler, T., Rust, J., Aberer, A. J., Aspöck, U., … Friedrich, F. (2014). Phylogenomics resolves the timing and pattern of insect evolution. Source: Science, 346(6210), 763–767. https://doi.org/10.2307/24917799

Morimoto, M. M., Nern, A., Zhao, A., Rogers, E. M., Wong, A. M., Isaacson, M. D., Bock, D. D., Rubin, G. M., & Reiser, M. B. (2020). Spatial readout of visual looming in the central brain of drosophila. ELife, 9, 1–102. https://doi.org/10.7554/ELIFE.57685

Muijres, F. T., Elzinga, M. J., Melis, J. M., & Dickinson, M. H. (2014). Flies evade looming targets by executing rapid visually directed banked turns. Science, 344(6180), 172–177. https://doi.org/10.1126/SCIENCE.1248955

Narasimha, S., Nagornov, K. O., Menin, L., Mucciolo, A., Rohwedder, A., Humbel, B. M., Stevens, M., Thum, A. S., Tsybin, Y. O., & Vijendravarma, R. K. (2019). Drosophila melanogaster cloak their eggs with pheromones, which prevents cannibalism. PLOS Biology, 17(1), e2006012. https://doi.org/10.1371/JOURNAL.PBIO.2006012

Oberst, S., Bann, G., Lai, J. C. S., & Evans, T. A. (2017). Cryptic termites avoid predatory ants by eavesdropping on vibrational cues from their footsteps. Ecology Letters, 20(2), 212–221. https://doi.org/10.1111/ELE.12727

Ohyama, T., Schneider-Mizell, C. M., Fetter, R. D., Aleman, J. V., Franconville, R., Rivera-Alba, M., Mensh, B. D., Branson, K. M., Simpson, J. H., Truman, J. W., Cardona, A., & Zlatic, M. (2015). A multilevel multimodal circuit enhances action selection in Drosophila. Nature, 520(7549), 633–639. https://doi.org/10.1038/nature14297

Parigi, A., Porter, C., Cermak, M., Pitchers, W. R., & Dworkin, I. (2019). The behavioral repertoire of Drosophila melanogaster in the presence of two predator species that differ in hunting mode. PLoS ONE, 14(5), 1–21. https://doi.org/10.1371/journal.pone.0216860

Pavan, M. (1948). Iridomyrmecin, an antibiotic substance extracted from the Argentine ant (Iridomyrmex pruinosus humilis Mayz.) VIII Internat. Cong. Entomol., Stock-Holm, 863–865.

Peters, R. S., Krogmann, L., Mayer, C., Donath, A., Gunkel, S., Meusemann, K., Kozlov, A., Podsiadlowski, L., Petersen, M., Lanfear, R., Diez, P. A., Heraty, J., Kjer, K. M., Klopfstein, S., Meier, R., Polidori, C., Schmitt, T., … Niehuis, O. (2017). Evolutionary History of the Hymenoptera. Current Biology, 27(7), 1013–1018. https://doi.org/10.1016/J.CUB.2017.01.027

Powell, J. R. (1997). Progress and prospects in evolutionary biology : the Drosophila model. Oxford University Press.

Reaume, C. J., & Sokolowski, M. B. (2006). The nature of Drosophila melanogaster. Current Biology, 16(16), R623–R628. https://doi.org/10.1016/J.CUB.2006.07.042

Refsnider, J. M., & Janzen, F. J. (2010). Putting Eggs in One Basket: Ecological and Evolutionary Hypotheses for Variation in Oviposition-Site Choice. Annual Review of Ecology, Evolution, and Systematics, 41, 39–57. https://doi.org/10.1146/ANNUREV-ECOLSYS-102209-144712

Ren, C., & Tao, Q. (2020). Neural Circuits Underlying Innate Fear. Advances in Experimental Medicine and Biology, 1284, 1–7. https://doi.org/10.1007/978-981-15-7086-5_1

Richmond, R. C., & Gerking, J. L. (1979). Oviposition site preference in Drosophila. Behavior Genetics, 9(3), 233–241. https://doi.org/10.1007/BF01066978

Rockwell, R. F., & Grossfield, J. (1978). Drosophila: Behavioral Cues for Oviposition. The American Midland Naturalist, 99(2), 361–368.

Rouquette, J., & Davis, A. J. (2003). Drosophila species (Diptera: Drosophilidae) oviposition patterns on fungi: The effect of allospecifics, substrate toughness, ovipositor structure and degree of specialisation. In Eur. J. Entomol (Vol. 100). https://doi.org/10.14411/eje.2003.056

Sawin-McCormack, E. P., Sokolowski, M. B., & Campos, A. R. (1995). Characterization and genetic analysis of Drosophila melanogaster photobehavior during larval development. Journal of Neurogenetics, 10(2), 119–135. https://doi.org/10.3109/01677069509083459

Schlenke, T. A., Morales, J., Govind, S., & Clark, A. G. (2007). Contrasting Infection Strategies in Generalist and Specialist Wasp Parasitoids of Drosophila melanogaster. PLoS Pathogens, 3(10), e158. https://doi.org/10.1371/journal.ppat.0030158

Sendoya, S. F., Freitas, A. V. L., & Oliveira, P. S. (2009). Egg-laying butterflies distinguish predaceous ants by sight. American Naturalist, 174(1), 134–140. https://doi.org/10.1086/599302

Sidhu, C. S., & Rankin, E. E. W. (2016). Honey Bees Avoiding Ant Harassment at Flowers Using Scent Cues. Environmental Entomology, 45(2), 420–426. https://doi.org/10.1093/EE/NVV230

Soto-Yéber, L., Soto-Ortiz, J., Godoy, P., & Godoy-Herrera, R. (2018). The behavior of adult Drosophila in the wild. PLoS ONE, 13(12), 1–26. https://doi.org/10.1371/journal.pone.0209917

Stensmyr, M. C., Dweck, H. K. M., Farhan, A., Ibba, I., Strutz, A., Mukunda, L., Linz, J., Grabe, V., Steck, K., Lavista-Llanos, S., Wicher, D., Sachse, S., Knaden, M., Becher, P. G., Seki, Y., & Hansson, B. S. (2012). A conserved dedicated olfactory circuit for detecting harmful microbes in Drosophila. Cell, 151(6), 1345–1357. https://doi.org/10.1016/j.cell.2012.09.046

Taylor, A. J., Muller, C. B., & Godfray, H. C. J. (1998). Effect of Aphid Predators on Oviposition Behavior of Aphid Parasitoids. Journal of Insect Behavior, 11(2), 297–302.

Traniello, J. (2010). Pheidole: Sociobiology of a Highly Diverse Genus. In Encyclopedia of Animal Behavior (Vol. 2, pp. 699–706).

Van Mele, P., Vayssieres, J. F., Adandonon, A., & Sinzogan, A. (2009). Ant cues affect the oviposition behaviour of fruit flies (Diptera: Tephritidae) in Africa. Physiological Entomology, 34(3), 256–261. https://doi.org/10.1111/j.1365-3032.2009.00685.x

Venken, K. J. T., Simpson, J. H., & Bellen, H. J. (2011). Genetic manipulation of genes and cells in the nervous system of the fruit fly. Neuron, 72(2), 202–230. https://doi.org/10.1016/j.neuron.2011.09.021

Vijendravarma, R. K., Narasimha, S., & Kawecki, T. J. (2013). Predatory cannibalism in Drosophila melanogaster larvae. Nature Communications 2013 4:1, 4(1), 1–8. https://doi.org/10.1038/ncomms2744

Wang, F., Wang, K., Forknall, N., Patrick, C., Yang, T., Parekh, R., Bock, D., & Dickson, B. J. (2020). Neural circuitry linking mating and egg laying in Drosophila females. Nature, 579(7797), 101–105. https://doi.org/10.1038/s41586-020-2055-9

Worthen, W. B., Natasha Hipp, M., & Twardokus, C. T. (1993). Effects of Ant Predation and Larval Density on Mycophagous Fly Communities. Oikos, 66(3), 526–532. Retrieved from https://www.jstor.org/stable/3544948

Yang, C., Belawat, P., Hafen, E., Jan, L. Y., & Jan, Y.-N. N. (2008). Drosophila egg-laying site selection as a system to study simple decision-making processes. Science, 319(5870), 1679–1683. https://doi.org/10.1126/science.1151842

Zhang, L., Yu, J., Guo, X., Wei, J., Liu, T., & Zhang, W. (2020). Parallel Mechanosensory Pathways Direct Oviposition Decision-Making in Drosophila. Current Biology, 30(16), 3075–3088.e4. https://doi.org/10.1016/j.cub.2020.05.076

